# Efficient transduction of pancreas tissue slices with genetically encoded calcium integrators

**DOI:** 10.1101/2025.03.21.644659

**Authors:** Charles S. Lazimi, Austin E. Stis, Julia K. Panzer, Helmut Hiller, Maria L. Beery, Amelia K. Linnemann, Cherie L. Stabler, Clayton E. Mathews, Edward A. Phelps

## Abstract

This study combines live pancreas tissue slices with viral transduction of the Calcium Modulated Photoactivatable Ratiometric Integrator 2 (CaMPARI2) biosensor for high-throughput analysis of islet calcium secretagogue responses. A key challenge of the pancreas slice model has been efficient transgene delivery throughout the slice volume while maintaining viability and function. Here, we demonstrate a robust adenoviral gene delivery approach to transduce slices with CaMPARI2 and apply photoconverting light to permanently mark glucose-induced calcium activity across all islets. This approach demonstrates glucose responsive CaMPARI2 labeling that correlates with insulin secretion. Using this novel high-throughput approach, we examine the relationship between islet size and calcium response. Larger isolated islets exhibit greater CaMPARI2 photoconversion in high glucose, whereas no size-function correlation is observed in islets resident in live slices. We also observe that slices capture a substantially higher proportion of small islets than isolated islets. Integrating CaMPARI2 with live pancreas slice studies enables multiplexed analyses, linking functional readouts to spatial features.

## Introduction

Pancreas tissue slices are an approach to study the physiology of islets and other pancreatic structures in the context of the native tissue microenvironment^1–8^. The slice technique maintains the architecture of the living tissue, including blood vessels, extracellular matrix, and immune cells. Slices avoid subjecting the pancreatic tissue to the harsh enzymatic and mechanical stress of whole islet isolation. Furthermore, slices allow for the investigation of crosstalk between different pancreatic tissue compartments^9^. Slices make it possible to monitor cell behavior in pathophysiological states that result in compromised islet morphology as well as in cases where isolation of whole islets is difficult, such as type 1 diabetes (T1D) and type 2 diabetes (T2D)^10^.

We and others have successfully used human and mouse pancreas slices to study pancreas physiology in non-diabetic donors and donors with T1D and T2D^10–16^. Functional studies in slices have mostly focused on Ca^2+^ imaging using chemical Ca^2+^ indicators, with some studies incorporating electrophysiology as well as observations of endocrine hormone secretion and exocrine enzyme release^3,8,10,17–19^. Two studies have used genetically encoded Ca^2+^ indicators (GECI) in slices generated from mice that express these transgenes^11,20^.

Robust protocols have been developed for viral transduction in isolated human pancreatic islets and reaggregated pseudo islets, facilitating the use of genetically encoded tools and biosensors^21–26^. Among many potential useful applications are GECI^27,28^, optogenetic and chemogenetic actuators^25,29–31^, gain or loss of gene function^32^, and ex vivo gene editing^33,34^. However, implementing viral gene delivery in human pancreas slices has been a challenge. A few examples of adenovirus transduction in slices have been reported. The Domínguez-Bendala group used adenovirus for lineage tracing^35,36^ and the Gaisano group demonstrated expression of LC3B-GFP^6^. However, these studies all showed low transduction efficiencies. Thus, consistent, robust viral transduction in pancreas slices has not yet been demonstrated.

Here, we report greatly improved adenoviral transduction in pancreatic tissue slices by introducing media agitation, physiological temperature culture, and removing the trypsin inhibitor. This approach substantially increased the number of transduced cells and facilitated the expression of genetically encoded biosensors while maintaining high tissue viability. Functional assays confirmed that transduced tissue slices preserved their responsiveness to stimuli, including real-time analysis of calcium signaling, oxidative stress, and insulin secretion.

We demonstrated the effectiveness of this transduction approach by implementing Calcium Modulated Photoactivatable Ratiometric Integrator 2 (CaMPARI2) to permanently encode Ca^2+^ responses in human pancreas slices. Current Ca^2+^ imaging approaches provide excellent temporal dynamics with real-time imaging of cell behavior including Ca^2+^ responses. However, such approaches are inherently low throughput, where the Ca^2+^ videos are captured one by one. It is typically only possible to obtain functional dynamics for a handful of islets per donor, which greatly under-samples the number of available islets in a slice preparation. Correlating live imaging with later immunostaining in the same islets is technically challenging. Here, we show that a photo-convertible Ca²⁺ integrator can be used to permanently encode Ca²⁺ responses in all islets simultaneously using slice preparations from donors without diabetes and from donors with T1D or T2D. The functional data from these tissues are preserved for correlation with fixed immunostaining. By linking these functional readouts with spatial parameters such as islet size, this method also enables exploration of how architectural features relate to glucose responsiveness. Together, this approach provides a valuable tool for investigating functional heterogeneity in the context of the in situ microenvironment and for studying disease mechanisms and therapeutic interventions.

## Methods

### Adenoviral generation

Plasmids, pAV-INS-GRX1-roGFP2 (VB240422-1276nau) and pAV-CMV-CaMPARI2 (VB240306-1476psj) were obtained from VectorBuilder for in-house adenovirus production. The plasmids were received as bacterial glycerol stocks and purified using the ZymoPURE II Plasmid Maxiprep Kit (Zymo Research, D4203-B). Plasmids were first linearized with the PacI restriction enzyme (New England Biolabs, R0547L) and precipitated using 100% isopropanol and 70% ethanol. The linearized DNA was transfected into HEK293T cells in a 6-well plate using Lipofectamine 2000 (Thermo Fisher Scientific, 11668027). Seven days post-transfection, HEK293T cells were examined for fluorescence and viral raft formation. By day 10, cells and media were collected, subjected to three freeze-thaw cycles using a dry ice/ethanol bath to lyse cells, and centrifuged to remove debris. 500 µl of viral supernatant was transferred to a fresh 10 cm plate of HEK293T cells for viral amplification. After two days, infected HEK293T cells underwent three freeze-thaw cycles for lysis. The adenovirus was subsequently harvested and purified using the Adenovirus Purification Kit (Cell Biolabs, VPK-5112). Adenovirus was then quantified using a QuickTiter Adenovirus Quantification kit (Cell Biolabs, VPK-106). Ad-INS-jRGECO1a and Ad-INS-GRX1-roGFP2 were provided by Dr. Linnemann at Indiana University. Ad-GCaMP6m (Vector Biolabs, 20190319T#9) was purchased from Vector Biolabs in ready-to-use high titer format.

### Slice culture and adenoviral transduction

Human pancreas tissue slices were generated by the Network for Pancreatic Organ Donors with Diabetes (nPOD) program at the University of Florida by embedding in agarose and cutting on a vibratome, according to published protocols^10,16,19^. Pancreatic tissue slices used for this study were 120 μm thick, and about 0.5 cm x 0.5 cm in length and width. For insulin secretion studies, pancreatic tissue (1 g) was obtained from Prodo Laboratories and shipped via courier to City of Hope, where slices were generated. After the slices were delivered by nPOD, they were allowed to recover for 24 hours in an incubator set to 24°C and 5% CO_2_ in a 10 cm petri dish with 10 ml of slice culture media consisting of Dulbecco’s Low Glucose Modified Eagles Medium +L-Glutamine +Pyruvate (Fisher Scientific SH30021.01), 10% fetal bovine serum (Sigma F4135), 25 kIU/ml aprotinin from bovine lung (Sigma A6106), and 1% antibiotic-antimycotic solution (Corning 30-004-CI). After letting the tissue slices recover for 24 hours, slices were transferred to 35 mm dishes at 2 slices per dish with 2 ml of slice culture media without protease inhibitor. For adenoviral transduction, 5.2 × 10^8^ PFU of ADV (2.6 × 10^8^ PFU per tissue slice) was added to each dish and slices were then cultured statically or on an orbital shaker (Vevor, 0-210) set to 40 rotations per minute inside a 37°C incubator. After 24 hours, viral-containing media was replaced with fresh human slice culture media without protease inhibitor and slices were cultured for an additional 24 hours on the orbital shaker at 37°C before imaging. For secretion studies under different culture conditions, Day 0 control was obtained on the day of slice generation after a 2h resting period on an orbital shaker at room temperature. Slices from Prodo Laboratories were cultured under the same conditions as nPOD slices.

### Perifusion

To evaluate glucose-stimulated insulin secretion, three viable pancreatic tissue slices were placed in a closed slice perifusion chamber (Biorep Technologies, PERI-PSC-PP and PERI-PSC-EXT) and connected to a perifusion system with automated tray handling (Biorep Technologies PERI5). The day 0 secretion control was obtained after 2 h resting on an orbital shaker at room temperature, other conditions were cultured for 24 or 48 hours in slice culture media with or without aprotinin and with or without the orbital shaker. Perifusion was performed with HEPES buffer (125 mM NaCl, 5.9 mM KCl, 2.56 mM CaCl2, 1 mM, MgCl2, 25mM HEPES, 0.1% BSA, pH 7.4) containing different glucose concentrations and 2 mM amino acids (L-alanine, L-arginine and L-glutamine (Sigma)). Slices were perfused at a flow rate of 100 μl/min. To remove accumulated hormones and enzymes, pancreatic slices were perfused with HEPES buffer containing 5.5 mM glucose for 90 minutes prior to perfusate collection. Slices were then sequentially exposed to different conditions: 5.5 mM glucose (5G) for 10 minutes and 16.7 mM glucose (16.7G) for 30 minutes. Perfusates were collected at a 60 sec interval in 96 well plates and stored at –20°C until insulin content was measured using a commercially available ELISA kit (Mercodia, human insulin ELISA, 10-1113-01).

### Calcium imaging

Live human pancreas tissue slices were transduced using the methods above with the Ca^2+^ biosensor Ad-CMV-GCaMP6m. Live imaging studies were conducted using a Leica SP8 confocal microscope equipped with a live cell incubation system allowing for long-term stable recordings with continual perfusion via an array of programmable syringe pumps (1000-US SyringeONE, New Era Pump Systems, Inc). Slices were initially perfused in Krebs-Ringer Bicarbonate Buffer with HEPES (KRBH, 115 mM NaCl, 4.7 mM KCl, 2.5LmM CaCl_2_, 1.2LmM KH_2_PO_4_, 1.2LmM MgSO_4_, 25LmM NaHCO_3_, 25 mM HEPES, 0.2% BSA) with 3 mM glucose (3G) then stimulated with KRBH with 16.7mM glucose (16.7G) and finally 3G with carbachol at 10 µM.

### Photoconversion of CaMPARI2

Human pancreatic tissue slices were received from nPOD and allowed to recover overnight at 24°C. Tissue slices were transduced with Ad-CMV-CaMPARI2 using the protocol listed above. After the 48-hour transduction period, slices were placed into an Ibidi 24-well plate (Ibidi 80821) with 750 µl of KRBH with 3 mM glucose at 37°C for 10 minutes to allow the slices to equilibrate. An additional 750 µl of 3 mM or 29 mM glucose was then added to the respective groups for 10 minutes before exposing the slices to 405 nm photoconverting (PC) light. Using a 405 nm light-emitting diode (LED) array (Amuza, LEDA-V), the PC light stimulus was delivered using 5 second pulses for 5 mins, leading to 2.5 minutes of total light delivery. The LED array evenly illuminated the entire 24-well plate with an intensity of 400 mW/cm^2^. An internal timer using an LED controller (Amuza, LAD-1) was used for precise timing of PC light exposure. For negative control groups, slices were incubated with 3 mM or 16 mM glucose but not exposed to PC light. Control groups were placed into a separate 37°C incubator to avoid any PC light cross-contamination. After PC light exposure, glucose solutions were aspirated, and slices were washed with PBS followed by overnight fixation with 4% paraformaldehyde (PFA) in PBS at 4°C.

### Immunohistochemistry

After an overnight fixation in 4% PFA, the slices were washed 3× with PBS and permeabilized with 0.1% Triton X-100 in PBS (TxPBS) for 30 minutes at room temperature (RT). Following permeabilization, the slices were blocked with 10% goat serum in TxPBS for 1 hour at RT to prevent non-specific antibody binding. Primary antibodies diluted in TxPBS with 1% goat serum were applied to the slices and incubated overnight at 4°C. The following day, the slices were washed three times for 5 minutes each with PBS at RT to remove unbound primary antibodies. Secondary antibodies and Hoechst 34580 were then applied and incubated for 1 hour at RT, followed by another three washes with PBS at RT. To prepare for mounting, a circle was drawn on a coverglass using a PAP pen liquid blocker (Newcomer Supply, NC9204359), and 200 µl of PBS was added to facilitate tissue transfer from the 24 well plate. The PBS around the tissue was then aspirated, and a drop of ProLong Gold antifade reagent (Invitrogen, P36930) was applied to the top of the tissue slice. A second coverslip was placed on top, utilizing a double coverslip technique with an imaging spacer (iSpacer, SunJin Lab IS211) to enable imaging from both sides of the tissue slice. The mounted slices were kept at RT overnight to allow the ProLong Gold to set, then stored at 4°C until imaging.

### Culture of human donor isolated islets and adenoviral transduction

Isolated and purified human islets were obtained from three non-diabetic donors from NIDDK-funded Integrated Islet Distribution Program (IIDP) at City of Hope. Human islets were removed from the shipping container, centrifuged at 180 x g for two minutes, and placed into Prodo Islet Media, PIM (S), (PRODO Labs PIM-S001GMP) supplemented with PIM(ABS) (Prodo Labs PIM-ABS001GMP), PIM(G) (Prodo Labs PIM-G001GMP), and PIM(3x) (Prodo Labs PIM-3X001GMP). Islets were separated into non-adherent cell culture dishes at a density of 1000 islet equivalents per dish. After a 24-hour recovery period at 24°C, islets were seeded into an 8-well ibidi dish (Ibidi 82421) at a density of 100 islets per well in 300 µl Prodo Islet Media. For adenoviral transduction, 1.6 × 10^7^ PFU of Ad-CMV-CaMPARI2 was added to each well and islets were then cultured inside at 37°C. After 24 hours, viral-containing media was replaced with fresh Prodo Islet Media and cultured for an additional 24 hours at 37°C before CaMPARI2 photoconversion and imaging. Photoconversion of isolated islets transduced with CaMPARI2 followed the same protocol used for slices.

### Tile scan analysis

Confocal imaging was performed using a Leica TCS SP8 confocal microscope equipped with a 20×/0.75 NA objective (Leica Microsystems). Tile scans of the entire pancreatic tissue slice were acquired to capture representative images of transduction efficiency throughout the whole tissue. A tile scan with a 10% overlap between adjacent tiles was used to ensure seamless stitching of the individual images. The scan area was defined manually, and the optimal number of tiles was automatically calculated by the Leica LAS X software. Pinhole size was set to 1 Airy unit for each channel, and the images were acquired at a resolution of 524 × 524 pixels per tile with a z-step size of 10 µm. For each field of view, a z-stack was acquired. The individual tiles were stitched into a single composite image using Leica LAS X software’s automated stitching algorithm. After stitching, images were processed using Fiji (ImageJ) software. Maximum intensity projections of the z-stacks were generated for further quantitative analysis.

### Statistics

All data are presented as mean ± SEM. One-way analysis of variance (ANOVA) followed by Tukey’s post hoc multiple comparisons test was applied to datasets presented in Figures 4, 5, and 6 to assess group differences. Outlier detection was conducted using the robust regression and outlier removal (ROUT) method with a false discovery rate (Q) set at 1%, identifying two outliers in the 16.7 mM glucose per islet condition for the T2D nPOD case 6619. Area under the curve (AUC) comparisons in Figures 1 and 2 were evaluated using an unpaired two-tailed t-test.

**Figure 1:**
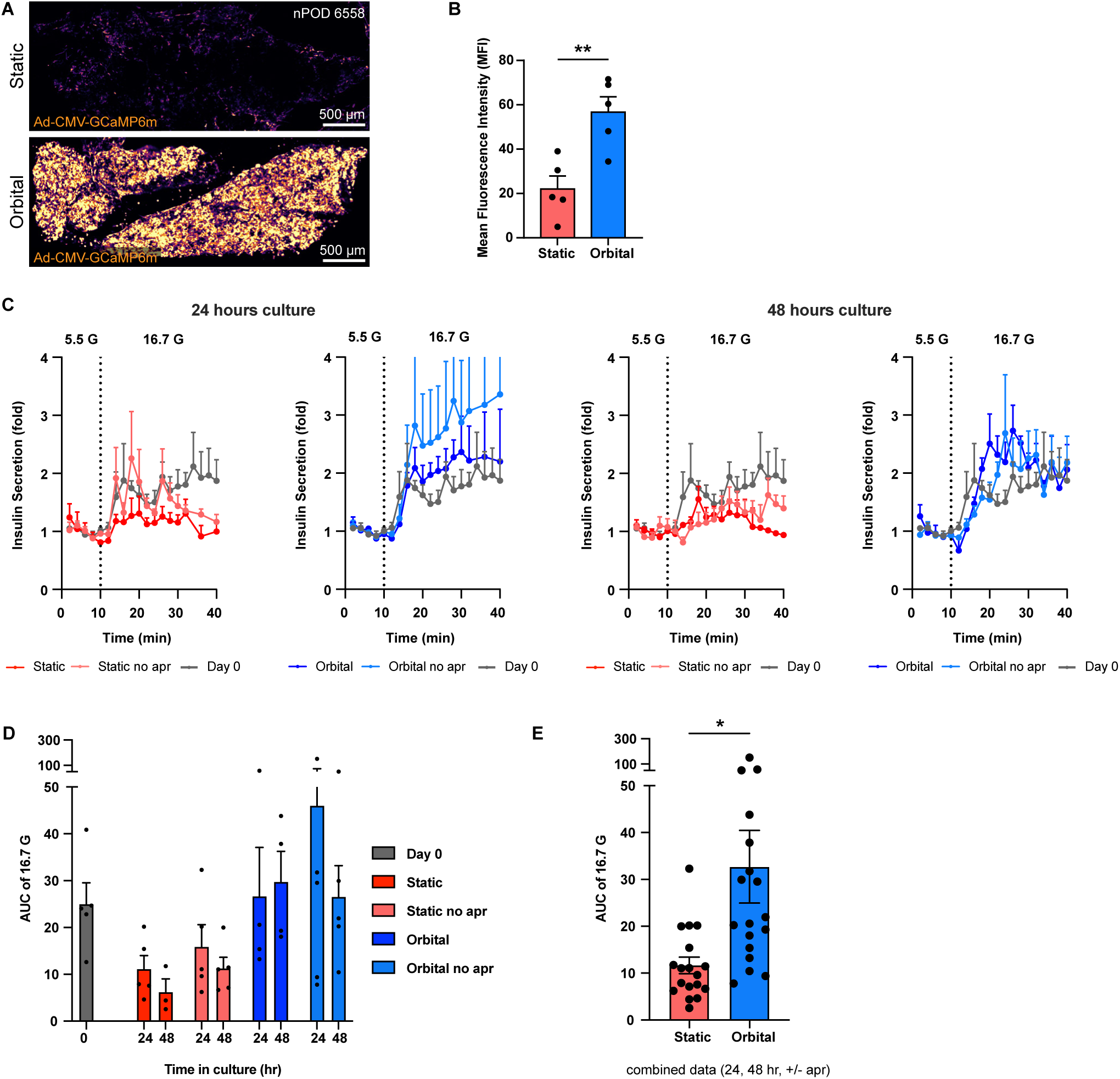
Orbital Shaker vs. Static Transduction and Insulin Secretion. **(A)** Confocal tilescan images of Ca²^+^ sensor Ad-CMV-GCaMP6m expression in whole human pancreatic tissue slices (nPOD 6558), transduced on an orbital shaker or in static culture with no aprotinin. Slices were transduced using 2.6 × 10L PFU of Ad-CMV-GCaMP6m per tissue slice, incubated for 24 hours at 37°C, and imaged 48 hours post-transduction. **(B)** Transduction efficiency was quantified using Mean Fluorescence Intensity (MFI), (± SEM, n = 5, students unpaired t-test, ** p < 0.01). **(C)** Dynamic insulin secretion of human pancreatic tissue slices from three to four nondiabetic donors were cultured on an orbital shaker (blue) or in static culture (red) after 24 and 48 hours, with and without aprotinin (apr). Insulin secretion traces from slices on Day 0 (gray) is repeated on all plots as a control. Slices were initially perfused with 5.5 mM glucose (5G) for 10 minutes, followed by stimulation with 16.7 mM glucose (16.7G) for 30 minutes. Data are normalized to the average baseline secretion at 5.5G. **(D)** Quantification of insulin secretion from perifusion assays in (C), shown as the area under the curve (AUC) for 16.7G stimulation at 0, 24 and 48 hours with or without aprotinin. AUC data from slices on Day 0 (gray) are provided as a control. **(E)** Same AUC data as in (D), but with aprotinin and no-aprotinin groups combined at 24 and 48 hours. (± SEM, n = 18, students unpaired t-test, * p < 0.05). The data are representative of the average of three to four donors, with each donor contributing four slices.

**Figure 2:**
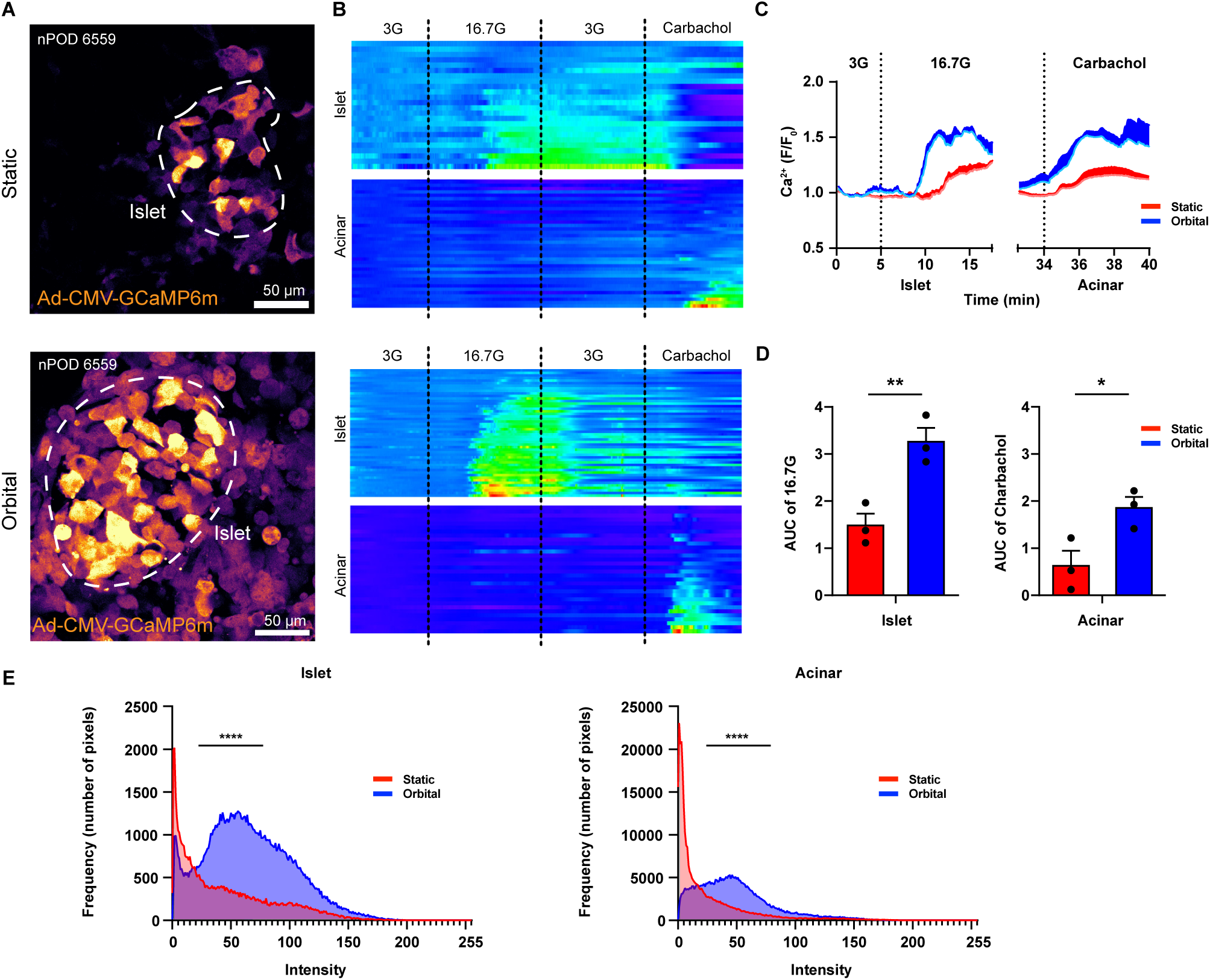
Endocrine and Exocrine Function Retained 72 Hours Post Adenoviral Transduction. Realtime confocal time-lapse perfusion study of pancreatic tissue slices (nPOD 6559) expressing the adenoviral Ca^2+^ biosensor CMV-GCaMP6m, transduced using an orbital shaker (bottom row) or static culture (top row). Endocrine and exocrine tissues were stimulated using perfusion of either 16.7 mM high glucose (16.7G) or 10 μM carbachol, respectively. **(A)** Images were selected from the real-time perfusion study at the peak Ca^2+^ influx of the 16.7 mM stimulation, islets outlined in white. **(B)** Heat maps of realtime Ca^2+^ responses in individual cells in both islet and acinar tissues shown in (A), imaged 48 hours post-transduction. Baseline activity was first acquired by perfusion of low 3 mM glucose (3G) for five minutes, followed by 16.7G, eliciting Ca^2+^ influx within the pancreatic islet. High glucose was washed away by perfusion of 3G, followed by stimulation of acinar tissue using 10 μM carbachol. **(C)** Average real-time Ca^2+^ traces of (n=3) slices following the protocol outlined in (B). The X-axis is segmented to separately show Ca^2+^ influx in the islet (left) and acinar tissues (right). **(D)** Quantification of the Ca^2+^ traces in (C), shown as the area under the curve (AUC) for 16.7G stimulation in the islet or 10 μM carbachol stimulation in the acinar tissue. (± SEM, n = 5, students unpaired t-test, * p < 0.05, ** p < 0.01) **(E)** Histograms depicting the sum of pixel intensity values at peak Ca^2+^ influx in the islet during high glucose stimulation and in the acinar tissue during carbachol stimulation. (Kolmogorov-Smirnov nonparametric cumulative distributions test, **** p <.0001).

**Figure 3:**
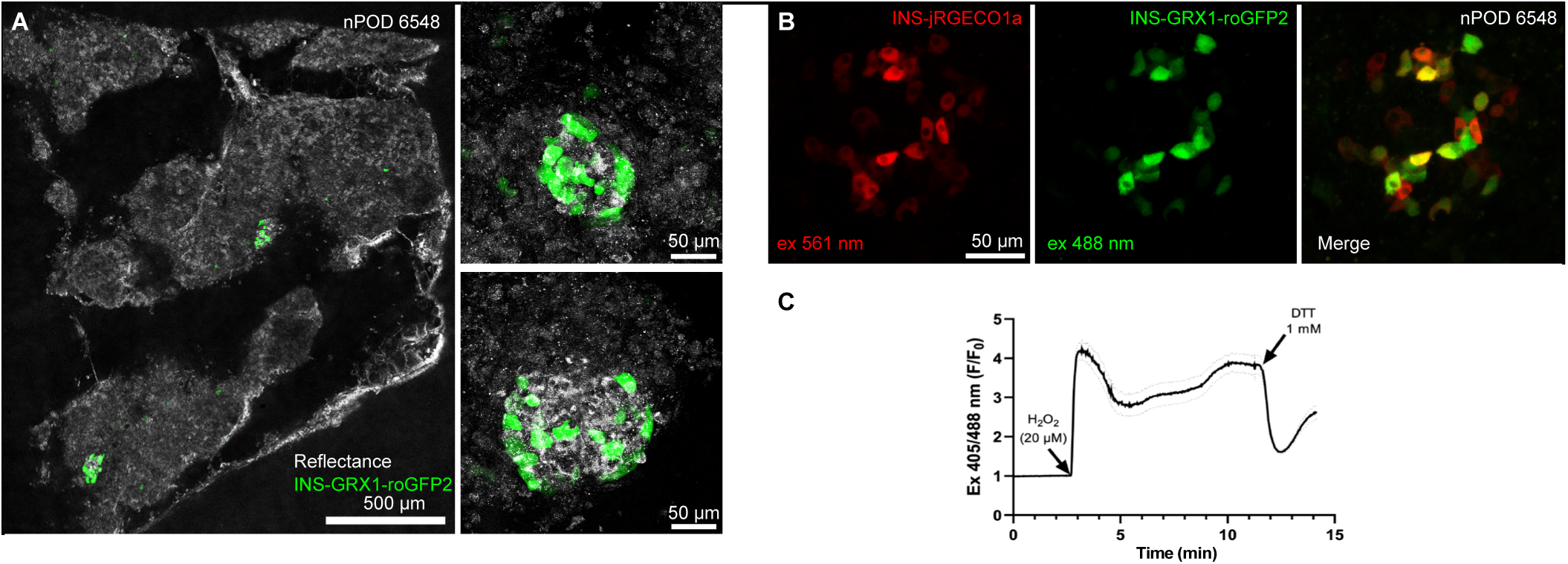
Targeted expression in specific cell types. Confocal images of beta cell-targeted expression of adenoviral vectors INS-GRX1-roGFP2 (green) and INS-jRGECO1a (red). **(A)** Representative image of a human pancreas slice alongside higher magnification images of islets within the tissue (nPOD 6548). Slices were transduced using 2.6 × 10L PFU of INS-GRX1-roGFP2 per tissue slice, incubated for 24 hours at 37°C, and imaged 48 hours post-transduction. **(B)** Dual expression of INS-GRX1-roGFP2 (green) and INS-jRGECO1a (red) in an islet within a human pancreas slice. Dual expression was achieved in slices using 1.4 × 10L PFU of INS-GRX1-roGFP2 and 1.4 × 10L PFU of INS-jRGECO1a per tissue slice, incubated for 24 hours at 37°C. Confocal images were taken 48 hours post-transduction. **(C)** Perfusion of human pancreatic tissue slice in a Warner imaging chamber using 3 mM glucose for 3 minutes, followed by a 20 μM H2O2 stimulation. ROS detoxification was achieved by perfusing 1 mM DTT at 12 minutes.

**Figure 4:**
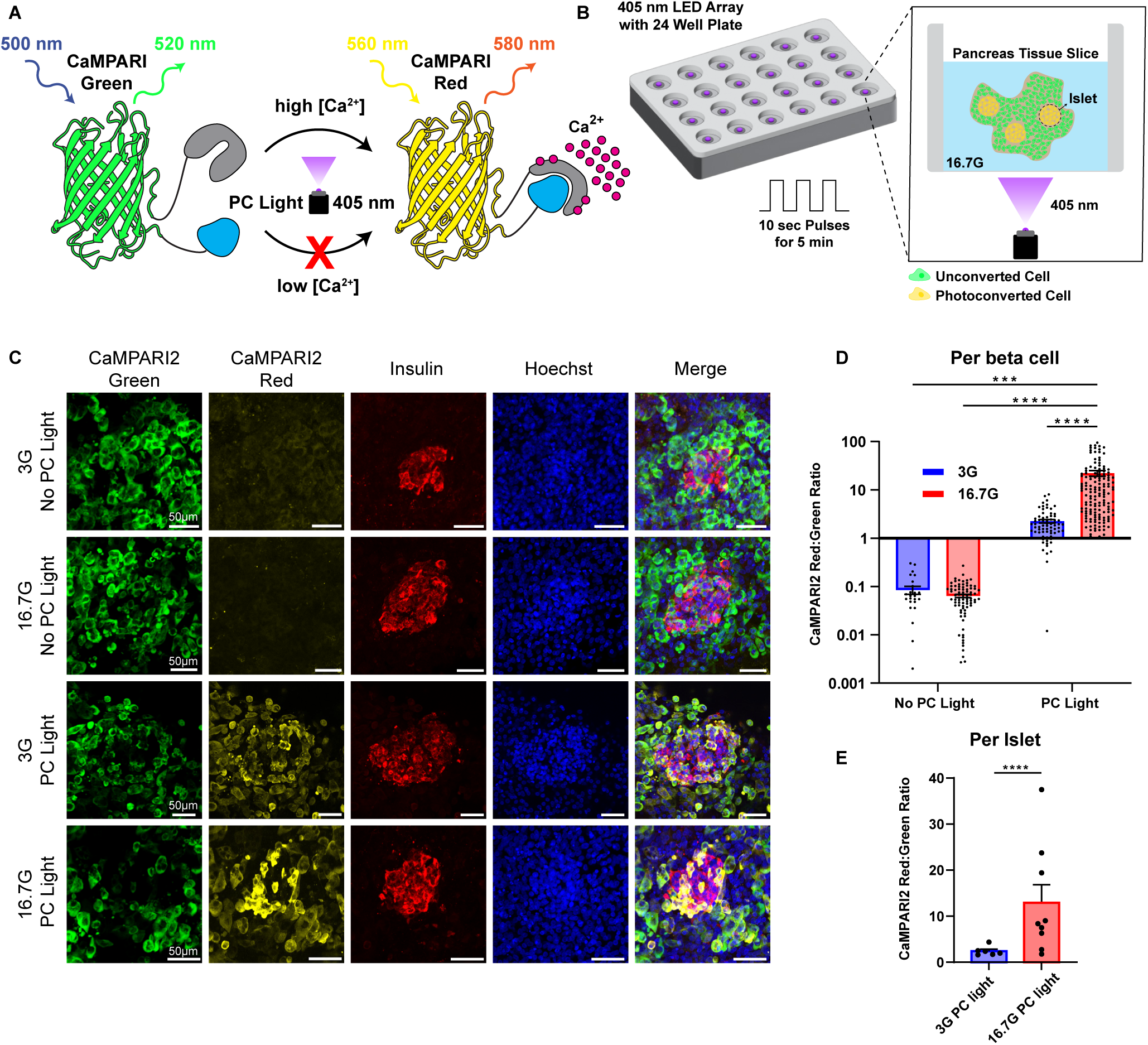
Functional Ca²+ Biosensor, CaMPARI2, in Living Human Pancreas Tissue Slices (nPOD 6617 ND). Cells that respond to glucose stimulation experience an influx of calcium ions. **(A)** An increase in intracellular calcium, along with exposure to ultraviolet (UV) light for photoconversion (PC), irreversibly converts the fluorescence of CaMPARI2 from green to red. **(B)** Schematic of CaMPARI2 setup including 24 well plate on 405 nm LED array controlled by pulse generator delivering 10 sec pulses for 5 min. (Inset) Single well containing transduced pancreatic tissue slice in 16.7G high glucose with photoconverted beta cells. **(C)** Slices transduced with the CaMPARI2 biosensor were incubated at 37°C in high or low glucose for ten minutes, followed by exposure or no exposure to PC light. PC light-treated groups were exposed using ten second pulses for five minutes. Tissue slices were then chemically fixed with paraformaldehyde for staining. **(D, E)** Glucose responsiveness in beta cells was quantified by calculating the CaMPARI2 red:green ratio. Only cells that were positive for insulin were quantified (n = 3 slices per group). Data were analyzed per beta cell or per islet. (± SEM, one-way ANOVA with Tukey’s post-hoc test, *** p < 0.001, **** p < 0.0001, outliers removed with ROUT Q = 1%).

Kolmogorov-Smirnov test was used to compare the cumulative distributions in the histograms of Figures 1 and 2. Slice and isolated islet CamPARI2 response vs area comparisons in Figure 8D were evaluated using an unpaired two-tailed t-test.

### Reagents

The following primary and secondary antibodies were used for immunofluorescence staining: Insulin Polyclonal Guinea Pig Antibody (Dako / Agilent no. A0564), Anti-CaMPARI Red 4F6 Mouse Antibody (Absolute Antibody no. AB01649), Goat anti-Mouse IgG (H+L) Highly Cross-Adsorbed Secondary Antibody Alexa Fluor™ 568 (Thermo Fisher no. A11031), Goat anti-Guinea Pig IgG (H+L) Highly Cross-Adsorbed Secondary Antibody Alexa Fluor™ 647 (Thermo Fisher no. A21450).

The following chemicals and reagents were used:

Carbachol (Thermo Fisher, AAL0667403), Dithiothreitol (Sigma, D9779), Hoechst 34580 (Invitrogen, H21486), Isopropanol (Fisher, A416-4), 200 Proof Ethanol (Decon, 2701), TE Buffer (Fisher, BP2473100), FBS (Gibco, 10082147), Sodium bicarbonate (Sigma, S5761), Sodium chloride (Sigma, S5886), PBS (Thermo Fisher Scientific, 20012050), HEPES (Sigma, H4034), HBSS (Gibco, 14025092), Penicillin Streptomycin (Gibco, 15140122), Potassium phosphate monobasic (Sigma, P5655), Magnesium sulfate heptahydrate (Sigma, M2773), BSA (Fisher Scientific, 199898), Calcium chloride (Sigma, C5670), Potassium chloride (Sigma, P5405), D-(+)-Glucose (Sigma, G7021), Paraformaldehyde (Sigma, P6148), Tolbutamide (Sigma, T0891), Diazoxide (Sigma, D9035), Forskolin (Sigma, F6886).

### Study approval

All experimental protocols using mice were approved by the University of Florida Animal Care and Use Committee (201808642, 202109991, 202400000147). All procedures using human slices were performed according to the established standard operating procedures of the nPOD/OPPC and approved by the University of Florida Institutional Review Board (IRB201600029) and the United Network for Organ Sharing (UNOS) according to federal guidelines, with informed consent obtained from each donor’s legal representative. Pancreas tissue from human donors with or without T1D was procured from the nPOD program at the University of Florida (RRID: <u>SCR_014641</u>, https://www.jdrfnpod.org). For each donor, a medical chart review was performed, and C-peptide was measured, with T1D diagnosed according to the guidelines established by the American Diabetes Association (ADA). Demographic data, hospitalization duration, and organ transport time were obtained from hospital records. Donor pancreata were recovered, placed in transport media on ice, and shipped via organ courier to the University of Florida. The tissue was processed by a licensed pathology assistant. Detailed donor information is listed in Table 1. nPOD tissues specifically used for this project were approved as nonhuman by the University of Florida IRB (NH00041892, NH00042022).

**Table 1:**
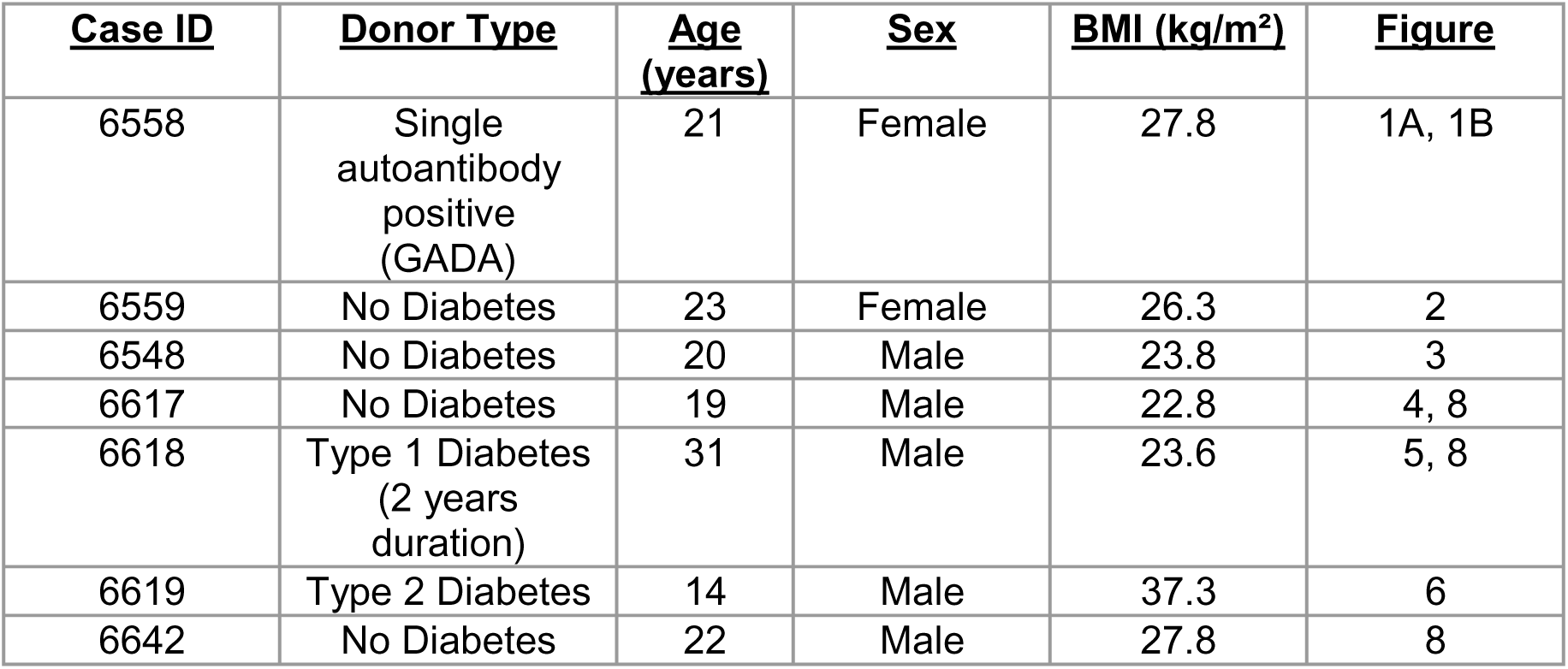
Donor characteristics from nPOD cases. Table summarizes donor information and applied figure.

**Table 2:**
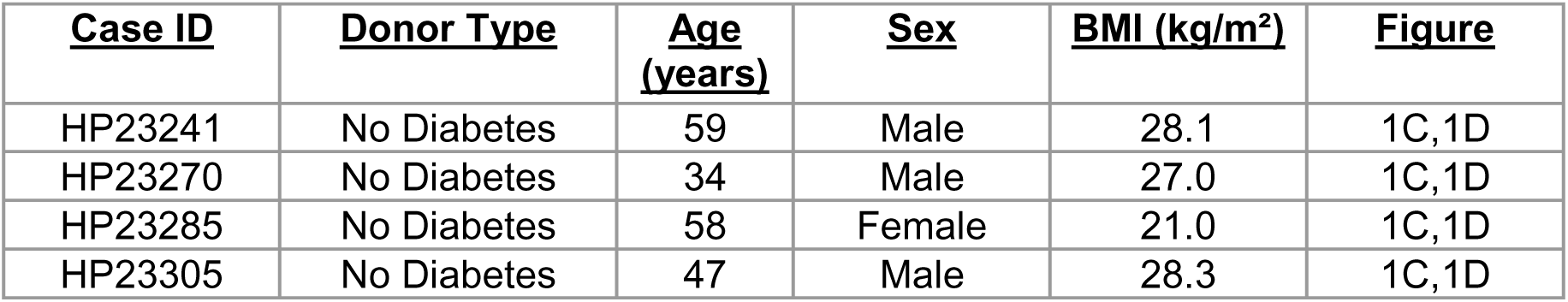
Donor characteristics from Prodo Labs. Table summarizes donor information and applied figure.

## Results

### Transduction on orbital shaker improves transduction efficiency

Robust expression of genetically encoded tools and biosensors for studying cellular processes has been challenging to achieve in pancreatic tissue slices. Here, we modified pancreas slice culture procedures to develop an approach for efficient viral gene delivery. Pancreas tissue slices secrete the digestive enzymes amylase, lipase, and trypsinogen, which can compromise slice viability^19^. To mitigate pancreatic enzyme activity, human pancreas slices are typically cultured with aprotinin at 28-30°C^10,16,19^. The lower culture temperature, combined with the use of trypsin inhibitor, prevents self-digestion and slows metabolism, thereby extending slice viability. These culture conditions also seem to hinder efficient viral transduction^37^. Additionally, in a static culture environment, transduction is limited by reduced viral particle collisions, as large particles like virions diffuse more slowly than small solutes^38^. We hypothesized that introducing media agitation would enhance viral mass transport to the tissue slices. Unlike monolayer cultures, where cells are evenly distributed, cells in tissue slices are highly concentrated in a small area. We hypothesized that agitation would help disperse the secreted pancreatic enzymes more efficiently than diffusion alone. This elevated efficiency should then allow for slice culture at 37°C without a protease inhibitor during the brief viral transduction period.

To test this hypothesis, we initially focused on delivering genetically encoded calcium indicators (GECI) to slices, as these biosensors allow us to assess islet cell function post-transduction by Ca^2+^ imaging. To improve pancreas tissue slice transduction, we removed protease inhibitor from the media, increased the culture temperature to 37°C, introduced media agitation by orbital shaker, and used a very high viral particle concentration of 5.2 x 10^8^ PFU/ml. A virus concentration of 1 x 10^6^ PFU/ml is typical for transducing cell monolayers. A standard viral titer for transducing cultured cells appears to be insufficient for good transduction of organotypic slices. Previous studies using adenovirus to transduce pancreas slices reported an MOI of 10 to 50 ^35,36,39^. We used an MOI of 86 in our protocol, assuming a typical pancreas slice is 5,000 µm x 5,000 µm x 120 µm in dimension and contains around 3×10^6^ cells, with two slices per dish in 2 ml of transduction media.

We initially compared the transduction of pancreas slices with and without media agitation, keeping MOI, lack of aprotinin, and temperature the same in both groups. At 24 hours, media was exchanged for fresh virus-free media, and we waited an additional 24 hours for transgene expression to occur. Transduction efficiency was quantified by taking confocal tile scans of the entire tissue slice and analyzing the mean fluorescence intensity (MFI) of the maximum intensity projection of each tissue slice (n=5 tissue slices per group) (Figure 1A). We observed low fluorescence and sporadic transduction in the statically cultured slices but widespread and bright fluorescence in the slices with media agitation. The simple addition of the low-speed orbital shaker during culture increased the mean fluorescence intensity of GCaMP6m by approximately 3x (Figure 1B).

While higher temperature culture, removal of protease inhibitor, and media agitation using an orbital shaker greatly improved transduction efficiency, these conditions could potentially be detrimental to islet function. To investigate any detrimental effects, we conducted a dynamic perifusion assay to evaluate insulin secretion from pancreatic tissue slices cultured under orbital shaker conditions compared to static culture after 24 and 48 hours and with and without the protease inhibitor aprotinin (apr) (Figure 1C). As measured by the area under the curve (AUC) during the stimulation phase, slices cultured on the orbital shaker at 37°C for 24 or 48 hours and with or without aprotinin were not statistically different from slices tested on the day of generation (day 0) or from slices cultured under static conditions with or without protease inhibitor (Figure 1D). These data indicate that the addition of orbital culture and removal of protease inhibitor were not detrimental to glucose responsive insulin secretion in slices. As human slices are a limited resource with substantial variability, we pooled the AUC datapoints from the different static and orbital groups to increase the sample size and compared static versus orbital head-to-head. In this comparison, orbital culture resulted in significantly improved insulin secretion over static culture (Figure 1E).

### Endocrine and exocrine function retained post adenoviral transduction

Adenoviral transduction in dense tissues often requires high multiplicities of infection (MOIs), which can result in cytotoxicity. While the tissue may express the biosensor, cellular responsiveness may be compromised. We demonstrate this by using the Ca^2+^ biosensor Ad-CMV-GCaMP6m, which allows for high transduction efficiency while maintaining both endocrine and exocrine cell responsiveness 72 hours post-adenoviral transduction when cultured on an orbital shaker. Human pancreas slices from a donor without diabetes transduced with the GECI Ad-CMV-GCaMP6m under either orbital or static conditions were assessed for dynamic Ca^2+^ responses to 16.7 mM glucose and the cholinergic agonist carbachol. Representative snapshots of peak Ca^2+^ influx are shown in Figure 2A, with heatmaps and dynamic perfusion traces of calcium responses presented in Figures 2B and 2C. Of note, fewer transduced cells were visible in the static culture group due to lower transduction efficiency and expression levels. Slices transduced under orbital shaker conditions had a significantly greater area of cells expressing the probe and brighter probe fluorescence within transduced cells (Figure 2E). Tissue slices transduced under orbital shaking conditions exhibited greater calcium influx in response to 16.7 mM glucose in endocrine cells and to 10 µM carbachol in exocrine cells, compared to slices transduced in static culture (n=3) (Figure 2D, Supplementary Video 1).

### Targeted expression in specific cell types

An advantage of genetically expressed biosensors over chemical sensors is the ability to target specific cell types using cell type specific promoters. In pancreatic tissue slices, where multiple cell types are present, insulin promoters ensure that data is collected only from beta cells. Here, we demonstrate that we can target beta cells using a reactive oxygen species (ROS) sensor driven by a hybrid insulin promoter (RIP / rabbit BGI)^40^, Ad-INS-GRX1-roGFP2 with minimal leaky expression (Figure 3A). We also show the ability to co-express multiple targeted sensors for multiplexed assays, using both a Ca^2+^ biosensor, Ad-INS-jRGECO1a, and Ad-INS-GRX1-roGFP2 (Figure 3B). GRX1-roGFP2 is a fusion of human glutaredoxin-1 (GRX1) to redox-sensitive GFP (roGFP2) that facilitates live imaging of the glutathione redox potential (E(GSH)) with high temporal resolution^40,41^. jRGECO is a red fluorescent protein-based GECI^42^. Following transduction, the function of the ROS probe and tissue responsiveness was confirmed by perfusing pancreatic tissue slices with hydrogen peroxide (H_2_O_2_). Upon 20 µM H_2_O_2_ exposure, an increase in the oxidation ratio of Ad-INS-GRX1-roGFP2 was observed, indicating successful detection of ROS. To further confirm the functionality and reversibility of the sensor, the tissue slices were subsequently treated with 1 mM dithiothreitol (DTT), a reducing agent (Figure 3C). This treatment successfully restored the baseline oxidation ratio, demonstrating that the ROS sensor retained its sensitivity and reversibility post-transduction. These data demonstrate the ability to specifically transduce beta cells in pancreas slices and multiplex multiple probes in a single slice.

### High throughput screening of beta cell glucose responsiveness in tissue slices

Calcium Modulated Photoactivatable Ratiometric Integrator 2 (CaMPARI2) is a photoconvertible protein that enables permanent marking of Ca^2+^ activity in large populations of cells over defined time windows. CaMPARI2 shifts from green to red emission upon simultaneous irradiation with violet light (405 nm) in the presence of elevated intracellular calcium (Figure 4A) ^43,44^. CaMPARI2 enables the permanent marking of cells that have experienced an increase in intracellular Ca^2+^ during a defined time window of photoconverting (PC) light exposure. Simultaneous PC light and elevated Ca^2+^ are required for photoconversion of the probe.

Real-time calcium imaging in human pancreatic tissues has traditionally relied on chemical sensors, such as Fluo-4, confined to the microscope’s live field of view. Here, we demonstrate the use of adenoviral transduction of pancreatic tissue slices with a genetically encoded Ca^2+^ integrator, Ad-CMV-CaMPARI2, allowing for high throughput calcium activity assessment by encoding total calcium activity across the entire tissue during a defined stimulation window. This approach facilitated the simultaneous functional assessment of all the islets across up to 24 slices from a single nPOD slice case in one experiment (∼60 islets). No microscope was involved during the stimulation and photoconversion protocol. Calcium activity was only quantified after-the-fact in fixed and immunostained slices by measuring the photoconversion within insulin positive cells.

First, we assessed a slice donor with no diabetes to confirm the function of CaMPARI2 in our system. CaMPARI2 requires the simultaneous presence of elevated Ca^2+^ and PC light to convert from green to red states. Live slices transduced with Ad-CMV-CaMPARI2 using the orbital shaker method were placed in a 24 well plate (1 slice per well) and statically exposed to 3 mM or 16 mM glucose with or without PC light from a 405 nm light-emitting diode array positioned directly under the cell culture plate (Figure 4B). Following PC light exposure, slices were fixed and immunostained for insulin and with an antibody that specifically labels only the photoconverted red form of CaMPARI2 and not the unconverted green form of CaMPARI2^44^. We used this monoclonal antibody to strongly label any cells that had calcium activity during the stimulation protocol. Fixed, stained, and mounted slices were imaged by confocal at a time of the investigator’s choosing, which neutralized much of the uncertainty and randomness surrounding when nPOD slice donor cases arrived and allowed for careful and meticulous observations of all islets in the sample.

By confocal imaging of the fixed and immunostained slices, no photoconversion was observed in the CaMPARI2 expressing slices that were not exposed to PC light. There was no photoconversion for either 3 mM or 16 mM glucose without PC light, demonstrating no spontaneous photoconversion of CaMPARI2 (Figure 4C). Low amounts of photoconversion were seen in 3 mM glucose with PC light, reflecting basal islet calcium activity. In contrast, strong photoconversion was observed with the combination of 16 mM glucose and PC light, confirming that beta cells responded to glucose stimulation by mobilizing cytosolic calcium and this response was encoded and preserved in the fixed slices (Figure 4C). We quantified the CaMPARI2 photoconversion in beta cells by hand-drawing regions of interest (ROIs) over cells double-positive for CaMPARI2 and insulin and measuring the ratio of mean fluorescence intensity of CaMPARI2 red to CaMPARI2 green on a per beta cell and per islet basis (Figure 4D,E). Interestingly, we observed significant heterogeneity in beta cell responses to glucose at both the single beta cells and single islet level. Despite an overwhelmingly clear response to glucose, two of the nine islets from this non-diabetic donor did not respond to 16 mM glucose, suggesting CaMPARI2 can be used to investigate islet functional heterogeneity.

Having demonstrated that the probe is functional for encoding calcium activity in beta cells within human pancreatic tissue slices, we aimed to investigate whether we could use this probe to assess beta cell function in various diseased states. We compared beta cell function in a second donor without diabetes, a donor with T2D, and a donor with T1D. Donor characteristics are reported in Table 1. The donor without diabetes, (nPOD case 6622) showed strong glucose responsive Ca^2+^ mobilization when assessed by photoconversion ratio of the CaMPARI2 probe. The beta cells displayed strong calcium activity upon high glucose and KCl stimulation, resulting in significant CaMPARI2 photoconversion (Figure 5A, 5B), consistent with dynamic insulin secretion data from this case (Figure 5C).

**Figure 5:**
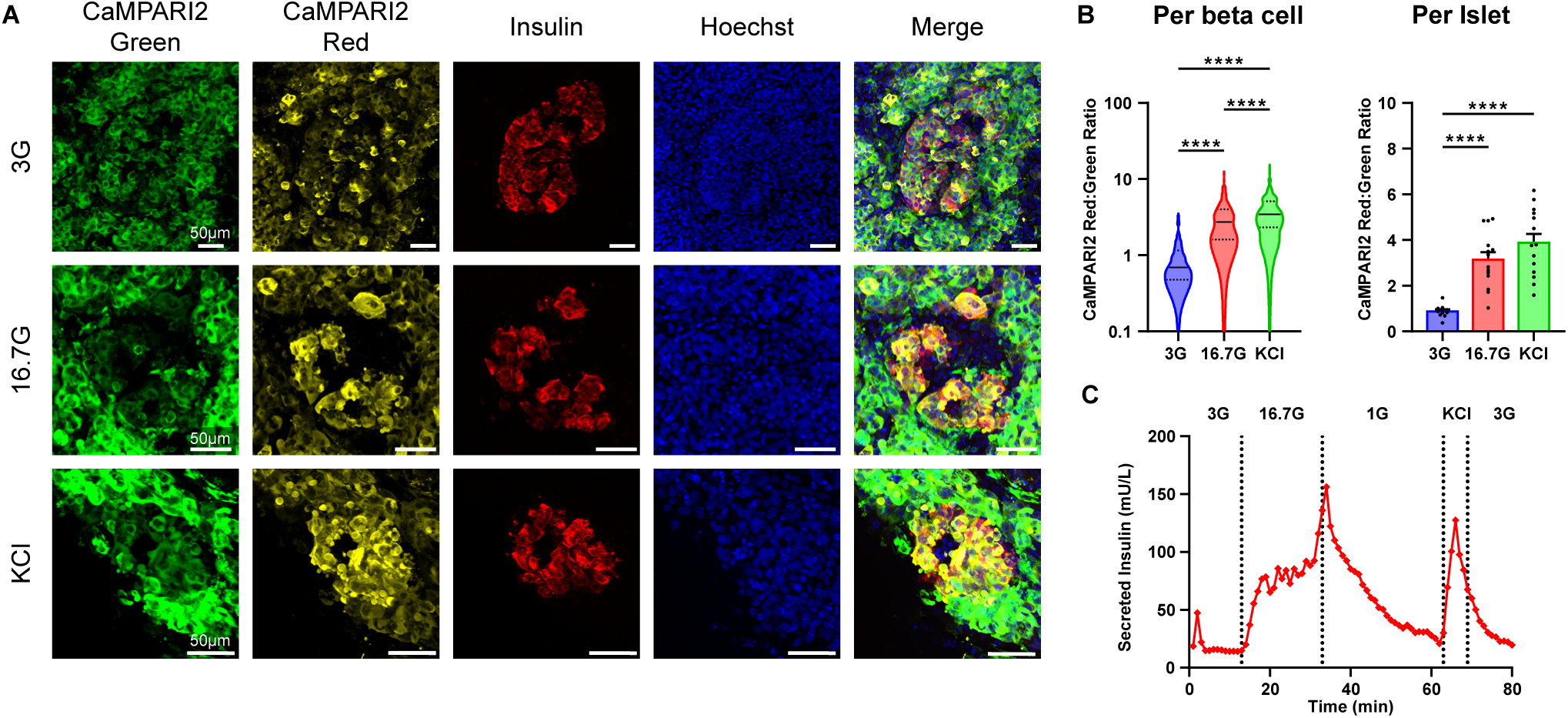
Functional Ca^2+^ Biosensor, CaMPARI2, in Living Human Pancreas Tissue Slices (nPOD 6622 ND Control). **(A)** Slices transduced with the CaMPARI2 biosensor were incubated at 37°C in 3 mM and 16.7 mM glucose for ten minutes, or 30 mM KCl for one minute. Slices were then exposed to PC light using ten second pulses for five minutes before being washed with PBS. Tissue slices were then chemically fixed with paraformaldehyde for staining. Beta cells displayed strong calcium activity upon high glucose stimulation, resulting in significant photoconversion. **(B)** Glucose responsiveness in beta cells was quantified by calculating the CaMPARI2 Red:Green Ratio. Only cells that were positive for insulin were quantified. (n = 4) slices per group. (± SEM, one-way ANOVA with Tukey’s post-hoc test, *** p < 0.001, **** p < 0.0001, outliers removed with ROUT Q = 1%). **(C)** Perifusion analysis of pancreatic tissue slices from this non-diabetic donor demonstrated robust insulin secretion in response to high glucose stimulation, confirming the preserved glucose responsiveness of beta cells.

Repeating the same experimental design for the calcium assay in a donor with T2D (nPOD case 6619), we observed a dysregulation in beta cell glucose sensing, particularly elevated CAMPARI2 conversion in low glucose conditions. (Figure 6A, 6B). This finding is consistent with insulin perifusion data, which also showed elevated insulin secretion at low glucose levels (Figure 6C). Additionally, the fold change in photoconversion between low and high glucose was markedly lower compared to the non-diabetic case, further highlighting the impaired glucose responsiveness in the beta cells of this particular T2D case, a phenotype that has been observed in other slices from donors with T2D^45^.

**Figure 6:**
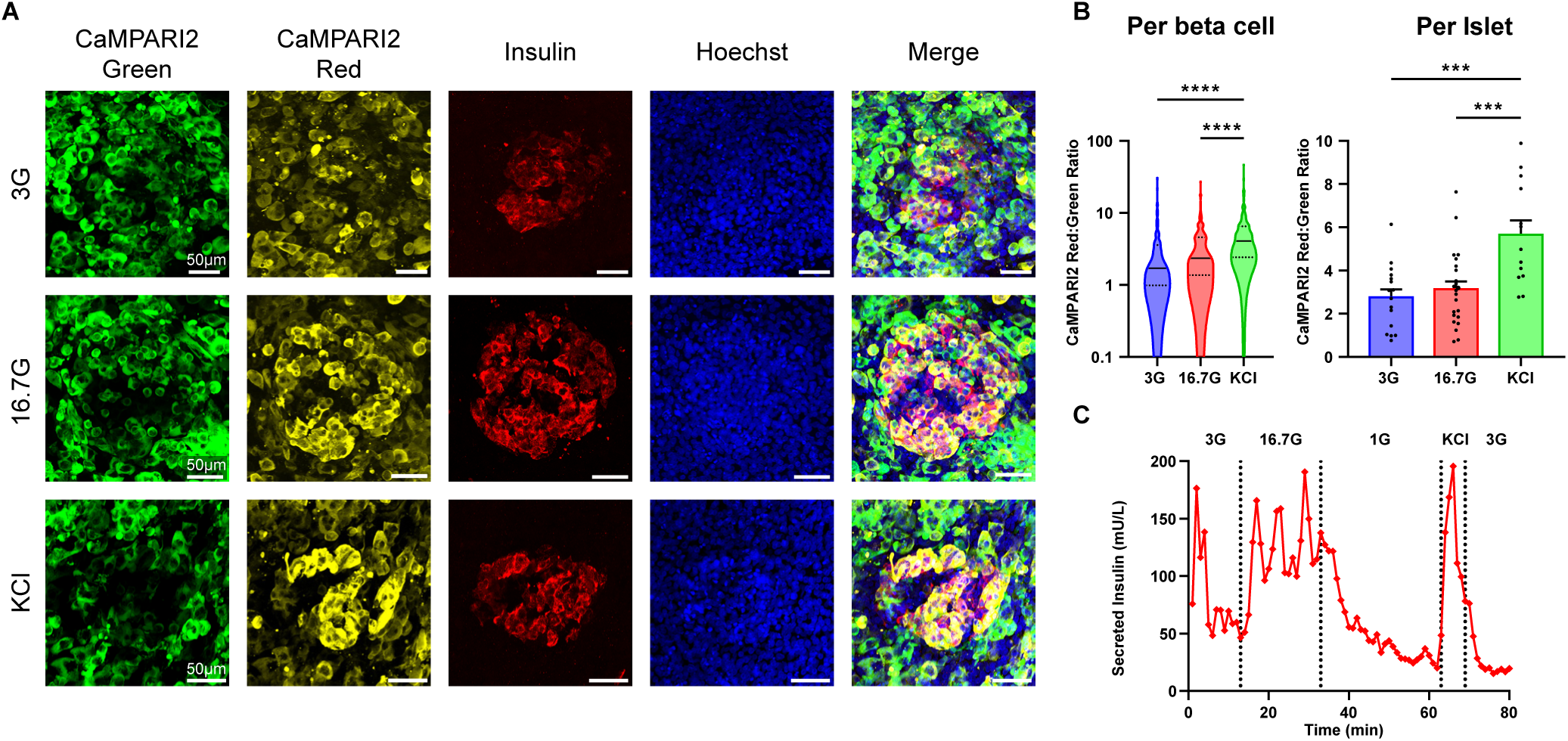
Functional Ca²^+^ Biosensor, CaMPARI2, in T2D nPOD Case 6619. **(A)** Slices transduced with the CaMPARI2 biosensor were incubated at 37°C in 3 mM and 16.7 mM glucose for ten minutes or 30 mM KCl for one minute. Slices were then exposed to photoconversion (PC) using ten second pulses for five minutes before washing with PBS. Tissue slices were then chemically fixed with paraformaldehyde for staining. **(B)** Glucose responsiveness in beta cells was quantified by calculating the CaMPARI2 Red:Green ratio. We observed a dysregulation in beta cell glucose sensing, particularly elevated calcium influx in low glucose conditions. Only insulin-positive cells were quantified (n = 4 slices per group). Data were analyzed on a per-beta-cell or per-islet basis. (± SEM, one-way ANOVA with Tukey’s post-hoc test, *** p < 0.001, **** p < 0.0001, outliers removed with ROUT Q = 1%). **(C)** The T2D case exhibited an atypical response, with significantly elevated insulin secretion at low glucose levels, aligning with the dysregulated calcium influx observed using CaMPARI2.

The CaMPARI2 assay was also performed in slices from a donor with T1D (nPOD case 6618), which had a disease duration of 2 years. Despite the significantly reduced beta cell mass in this case, we successfully captured the calcium activity from two remaining insulin-positive islets, one each under low and high glucose conditions. The residual beta cells were non-functional, showing no response to high glucose stimulation, resulting in low photoconversion of CaMPARI2 (Figure 7A, 7B). These findings are consistent with perifusion data for this case, which demonstrated a total absence of insulin release in response to both high glucose and KCl stimulations (Figure 7C). This data is also consistent with beta cell glucose blindness observed in live Ca^2+^ imaging in pancreas slices from donors with T1D^10,16,19^.

**Figure 7:**
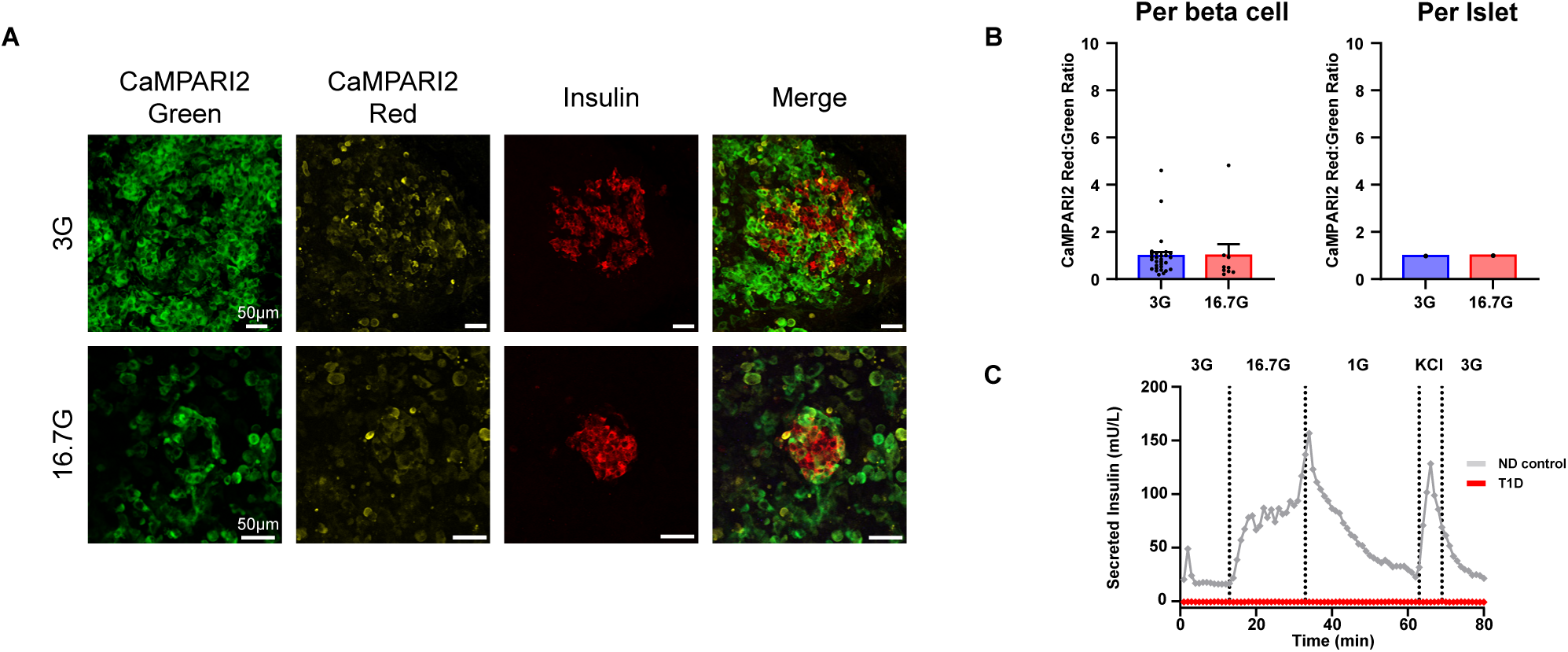
Functional Ca²^+^ Biosensor, CaMPARI2, in T1D nPOD case 6618. **(A)** Slices transduced with the CaMPARI2 biosensor were incubated at 37°C in 3 mM and 16.7 mM glucose for ten minutes. Slices were then exposed to PC light using ten second pulses for five minutes before washing with PBS. Tissue slices were then chemically fixed with paraformaldehyde for staining. **(B)** Glucose responsiveness in beta cells was quantified by calculating the CaMPARI2 red:green ratio. The residual beta cells were non-functional, showing no response to high glucose stimulation, resulting in no photoconversion of CaMPARI2. Only cells that were positive for insulin were quantified (n = 1 slice per group). Data were analyzed on a per-beta-cell basis. **(C)** Perifusion analysis of pancreatic tissue slices from this T1D case revealed a total absence of insulin release in response to both high glucose and KCl stimulations, confirming the non-functionality of the remaining beta cells (red). Perifusion trace from non-diabetic (ND) case 6622 (grey) provided as a control.

**Figure 8:**
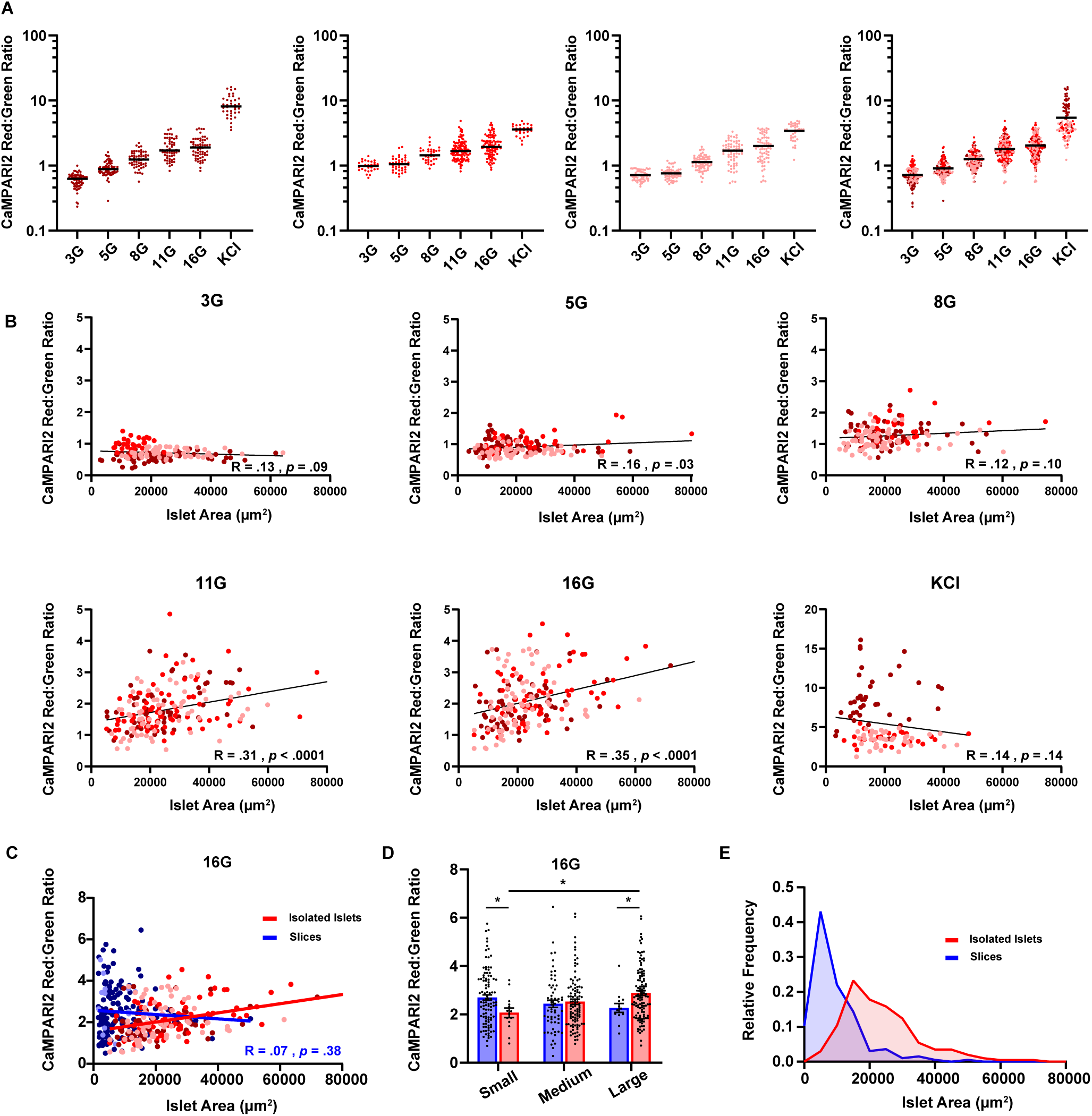
Functional Ca²+ Biosensor, CaMPARI2, Versus Islet Area in Human Isolated Islets and Slices (ND). **(A)** CaMPARI2 Red:Green ratio of human isolated islets in response to increasing concentrations of glucose and KCl. Plots are separated by donor (n = 3) indicated by the different shades of red, with each point representing a single islet. **(B)** CaMPARI2 response of the three isolated islet donors in (A), plotted against islet area (µm²) for each glucose concentration (3, 5, 8, 11, 16 mM), with each point representing a single islet. **(C)** CaMPARI2 Red:Green ratio at 16 mM glucose versus islet area (µm²) in both isolated islets (red) and slices (blue) (n=3; nPOD6617, nPOD6618, nPOD6642). **(D)** Isolated islets (red) and islets in slices (blue) were categorized as Small (<8,000 µm²), Medium (8,000–20,000 µm²), or Large (>20,000 µm²), and plotted to show differences in islet size and function (± SEM, n = 18, Student’s unpaired t-test, *p < 0.05). **(E)** Histogram showing the relative frequency distribution of analyzed islet areas in isolated islets (red) and slices (blue).

### Islet size correlates with Ca^2+^ responses in isolated islets but not slices

We used CaMPARI2 to assess whether islet area contributes to functional variability in human isolated islets and pancreatic tissue slices. We first assessed isolated islets from three donors without diabetes (Table 3). Using isolated islets, we assessed the Ca^2+^ response in ∼350 total islets per donor, ∼55 islets per glucose condition. The CaMPARI2 response showed stepwise increases in Ca²⁺ influx with progressively higher glucose (Figure 8A). To examine the size versus function relationship, we plotted each islet’s CaMPARI2 response against the measured area (µm^2^) for each glucose concentration (Figure 8B). At sub-activating glucose levels (3 and 5 mM), the slope of Ca²⁺ response versus islet area was flat, consistent with a lack of Ca^2+^ response at these glucose levels (correlation coefficient, R = 0.13 and 0.16; significance, p = 0.09 and 0.03). At 8 mM glucose, some islets started to respond, but there was no correlation with islet area (R = 0.12, p = 0.10). As glucose increased to activating levels (11 and 16 mM), Ca^2+^ responses were much stronger and the correlation between size and function became positive, indicating that larger isolated islets tended to exhibit greater Ca²⁺ responses under stimulatory glucose conditions (R = 0.31 and 0.35, p < 0.0001). At maximal stimulation by KCl, there was no size-function correlation (R = 0.15, p = 0.14).

**Table 3:** Isolated islet donor characteristics from Integrated Islet Distribution Program (IIDP) cases. Table summarizes donor information and applied figure.

| <u>Case ID</u> | <u>Donor Type</u> | <u>Age (years)</u> | <u>Sex</u> | <u>BMI (kg/m<sup>2</sup>)</u> | <u>Figure</u> |
| --- | --- | --- | --- | --- | --- |
| SAMN48357134 | No Diabetes | 37 | Female | 26.4 | 8 |
| SAMN48585017 | No Diabetes | 50 | Male | 27.9 | 8 |
| SAMN48756137 | No Diabetes | 35 | Female | 31.6 | 8 |

We next compared the size and function relationship between isolated islets and slices at 16 mM glucose from three donors each (Figure 8C). We did not observe a size-function correlation in slices at 16 mM glucose (R = 0.07, p = 0.38). Grouping islets into small (<8,000 μm², <100 μm diameter), medium (8,000–20,000 μm², 100-160 μm diameter), and large (>20,000 μm², >160 μm diameter) demonstrated that Ca²⁺ responses increased with size in isolated islets but remained relatively constant across size categories in slices (Figure 8D). Notably, small islets showed reduced CaMPARI2 responses in isolated islet preparations compared to slices. We quantified the islet size distributions in isolated islets and slices in our samples. Slices captured a substantially greater proportion of small islets than isolated preparations, which likely reflects loss of and damage to small islets during the digestion and isolation process (Figure 8E).

Overall, our findings demonstrate the utility of CaMPARI2 as a biosensor for permanently encoding calcium activity in human pancreatic tissue slices during a defined stimulation window. This tool can be implemented across various disease states, including T1D and T2D, as well as at different stages of disease progression, offering a powerful, high throughput means to assess glucose responsiveness and its relationship to islet architecture, including size, morphology, and location.

## Discussion

Adenoviral transduction in pancreatic tissue slices presents a unique challenge as the slices continuously secrete digestive enzymes^19^, which can degrade viral particles and impact the viability of the tissue. Additionally, as opposed to transduction of cell monolayers, where cells are evenly spread out across the culture dish, cells within a tissue slice are densely confined to a small area, reducing the probability of viral particle collisions. A study of adenoviral transduction of human brain slices used microinjection of high titer virus directly into the slice tissue to achieve sufficient transduction efficiency^46^. In static culture conditions commonly used for transduction, viral particles diffuse more slowly than smaller molecules, reducing their interactions with the tissue^38^. Our study demonstrates that gentle media agitation during adenoviral transduction significantly improves transduction efficiency compared to static conditions, without compromising tissue functionality. The increased transduction efficiency, evidenced by a 300% rise in MFI, suggests that media agitation facilitates the movement of viral particles, overcoming diffusion limitations and improving the chances of viral particles reaching the small surface area of pancreatic tissue slices. We anticipate that our approach will be conducive to transducing organotypic slices from different tissues and with other virus types including lentivirus and adeno-associated virus.

A potential limitation of orbital shaking is the introduction of mechanical stress, which could adversely affect beta cell function. To address this concern, we evaluated the functional integrity of transduced tissue slices through dynamic perifusion assays. Our results demonstrated that pancreas slices cultured under mild orbital shaking conditions not only retained but enhanced their insulin secretion capacity in response to glucose. This suggests that gentle orbital shaking does not negatively impact beta cell function; instead, it may contribute to maintaining tissue viability and responsiveness.

In addition to enhanced transduction efficiency, we showed that adenoviral transduction does not compromise the functional responsiveness of either endocrine or exocrine cells using real time calcium imaging. Calcium imaging using the genetically encoded calcium indicator Ad-CMV-GCaMP6m revealed strong calcium responses to glucose in islets and to carbachol in exocrine cells, further highlighting the preserved functionality of the transduced tissue. Tissue slices transduced under orbital shaking conditions exhibited a greater number of transduced cells and stronger calcium responses compared to static culture. A variety of probes can be used in slices, targeted to specific cell types, and even multiplexed, as shown by our demonstration of Grx1-roGFP2 and jRGECO1a.

The use of the genetically encoded calcium integrator CaMPARI2 allowed us to conduct high throughput calcium assays, facilitating the functional assessment of all the islets in an allocation of slices from an nPOD donor case (usually, 20 slices from each donor). The transient nature of traditional calcium indicators, such as Ad-CMV-GCaMP6m used in Figure 3, provide a limited field of view and require continuous monitoring of islets to capture peak calcium activity upon glucose stimulation. Selecting which islets to record can be subject to investigator bias, and data collected are not guaranteed to be representative of the entire islet population. All calcium recordings within a case are typically performed on the same day to minimize variability, but due to the time required to set up and run each recording, only 3 to 4 islets per case are usually captured. These constraints highlight the advantages of genetic tools like CaMPARI2 in pancreatic tissue slices. By enabling the simultaneous analysis of entire islet populations, CaMPARI2 reduces islet selection bias and ensures data capture from many more islets. This is especially critical in rare cases, such as donors with T1D, where samples may only be available sporadically and islet morphology is heterogeneous. Having a tool like CaMPARI2 ensures we maximize the value of the sample, increasing the likelihood of capturing representative data.

Efficient protocols for adenoviral transduction in slices enabled us to examine calcium dynamics across different disease states, including T1D and T2D cases. Our findings revealed dysregulated glucose sensing in T2D case 6619, characterized by elevated calcium influx under low glucose conditions, consistent with abnormal insulin secretion observed in perifusion assays. In T1D case 6618, we observed non-functional beta cells that failed to respond to high glucose stimulation, reflecting the loss of glucose-responsive insulin secretion in this case. While these findings are promising, they are based on a single case each. In this paper, we demonstrate the feasibility of using this tool to monitor calcium dynamics in various states, but further studies with more cases are needed to establish meaningful patterns or differences across disease states. Nonetheless, our work shows that CaMPARI2 is a powerful biosensor for detecting changes in calcium activity in beta cells within human pancreatic tissue slices across various disease stages.

Functional heterogeneity among islets may reflect differences in cellular composition, architecture, and size. In humans, both islet architecture and cellular composition are size-dependent^47–49^. This is particularly important because human islet sizes are highly heterogeneous, ranging from single endocrine cells to large structures composed of several thousand cells^47^. Several studies have reported that smaller islets are devoid of glucagon, whereas larger islets contain proportionally fewer insulin-positive cells^50–52^. Islet size has also been linked to susceptibility during disease progression, with smaller islets reported to be more vulnerable to autoimmune destruction in T1D^52–54^, while larger islets appear to be preferentially lost in T2D^47,49^. Here, we used CaMPARI2 to investigate the relationship between calcium responsiveness and islet size. In isolated islets, we observed a positive correlation between islet size and Ca²⁺ response, whereas no such correlation was detected in pancreatic tissue slices. These findings contrast with previous reports showing a negative correlation between islet size and insulin secretion, in which smaller islets secreted more insulin per volume than larger ones^51,55,56^. While Ca²⁺ influx is necessary for insulin secretion, many downstream processes can modulate the final amount of insulin released, meaning changes in Ca²⁺ entry do not necessarily equate one to one with insulin release.

Mean islet size is significantly smaller when measured by IHC in paraffin sections than for isolated islets^50^. We observed a similar trend in islet size for slices compared to isolated islets. In slices, 40% of analyzed islets were classified as small, compared to less than 10% in isolated preparations (Figure 8E). Data presented here, together with other reports, emphasize that isolated islets do not represent the full range of islet sizes present in the endogenous tissue^50,57^. In contrast, histological analysis of whole pancreas sections allows the study of the complete range of islets. However, this approach is limited to fixed tissue and cannot be integrated with multiplexed, live functional assays. Perifusion of live tissue slices also averages insulin secretion across several islets, obscuring size-dependent effects. Here, by combining CaMPARI2 with live pancreas slices, we present one of the first high throughput analyses comparing islet size to functional calcium dynamics in live human tissue.

Our findings also revealed significant heterogeneity in beta cell calcium responses within individual islets. This variation may reflect functional subpopulations of beta cells, such as first-responder or leader cells, which have been proposed to initiate coordinated islet activity. Studies have identified leader cells as highly responsive beta cells that drive islet-wide calcium waves^58–60^. Future studies using CaMPARI2 in conjunction with functional and transcriptomic profiling could help determine the identity of high or low responding cells and their role in glucose regulation, providing a molecular context to functional heterogeneity. Furthermore, CaMPARI2 can be combined with immune cell staining techniques to quantify islet function during insulitis, offering insights into how immune cell presence might impact beta cell function and survival. These capabilities open opportunities for understanding complex dynamics within pancreatic tissue slices, particularly in diseases like T1D, where both functional and immune-mediated factors are critical.

## Conclusions

Our study establishes an optimized adenoviral transduction protocol for human pancreatic tissue slices, significantly enhancing transgene delivery while preserving tissue viability and function. By incorporating media agitation, higher temperature culture, and removal of trypsin inhibitors during transduction, we significantly improved transduction efficiency without compromising endocrine or exocrine cell responsiveness. These optimizations address a critical technical limitation in pancreas slice research, facilitating the application of genetically encoded biosensors and functional genetic manipulations in human pancreatic tissue.

## Acknowledgements

This research was performed with the support of the Network for Pancreatic Organ donors with Diabetes (nPOD; RRID:SCR_014641), a collaborative type 1 diabetes research project supported by Breakthrough T1D and The Leona M. & Harry B. Helmsley Charitable Trust (Grant# 3-SRA-2023-1417-S-B). The content and views expressed are the responsibility of the authors and do not necessarily reflect the official view of nPOD. Organ Procurement Organizations (OPO) partnering with nPOD to provide research resources are listed at https://npod.org/for-partners/npod-partners/.

This work was funded by the following NIH grants: P01 AI042288 (EP), R01 DK132387 (EP), R01 DK124267 (EP), R01 DK123292 (EP), the NIDDK-supported Human Islet Research Network (HIRN, RRID:SCR_014393; https://hirnetwork.org) UH3 DK122638 (CLS, CEM, EP), and by Breakthrough T1D grant 2-SRA-2023-1313-S-B (EP).

The authors thank the nPOD donors and their families for the gift of tissues, Stephan Speier (Technische Universität Dresden) and Denise Drotar (University of Florida) for assistance in initiating our work in human pancreas slices, and Andrece Powell (University of Florida) for technical assistance with immunostaining.

## Author contributions

CSL developed the methodology for culturing and transducing live human pancreatic tissue slices on an orbital shaker, conducted fixed and real-time Ca²⁺ imaging studies, performed fixed tissue staining and imaging, analyzed Ca²⁺ imaging and CaMPARI2 data, generated adenovirus, and drafted the manuscript. AES helped design the setup for CaMPARI2 experiments, assisted with CaMPARI2 experiments, contributed to the development of imaging approaches, and generated adenovirus. JKP performed perifusion studies and insulin ELISAs on human tissue slices. AKL provided critical reagents and technical assistance. MB and HH generated live human pancreatic tissue slices and conducted perifusion studies. CLS and CEM provided critical funding, guidance and technical assistance. EAP secured funding, conceived the study, supervised the research, and edited the manuscript. All authors contributed to data interpretation, reviewed and commented on the manuscript, and approved its final submission.

